# Hepatic Differentiation of Human Pluripotent Stem Cells into Functional In Vitro Models Recapitulating Native Liver Complexity for MASLD Modelling

**DOI:** 10.64898/2026.06.02.729501

**Authors:** Saloni Sainger, Anshul Chikara, Manisha Kumari, Deepika Kumari, Pratibha Jaipal, Shubhanshi Ranjan, Sunil Gujjar, Yashwant Kumar, Ajay Kumar, Santosh Mathapati

## Abstract

Human *in vitro* hepatic models that accurately recapitulate liver function are essential for fundamental and translational research; however, currently utilised models for disease modelling and drug discovery lack physiological fidelity and require prolonged culture time. Here, we present a streamlined 10-day protocol for efficient and reproducible differentiation of human pluripotent stem cells into hepatocyte like cells (HLCs) and hepatic liver organoids (HLOs). Both models exhibited mature hepatocyte differentiation, as evidenced by albumin secretion and CYP3A4 metabolic activity. Interestingly, HLOs display enhanced multicellular complexity incorporating endothelial, stellate, and macrophage populations along with hepatocytes, thereby more closely recapitulating the native liver microenvironment than HLCs. Here, steatosis was induced in both platforms, which resulted in triglyceride accumulation and upregulated lipogenic markers (DGAT1, DGAT2). However, only HLOs recapitulated advanced disease characteristics, including inflammatory (IL-10) and fibrotic (alpha SMA, COL1A1) responses. Resmetirom, a thyroid hormone receptor-β agonist, significantly reduced steatosis and restored molecular signatures in both models. Additionally, transplanted HLOs demonstrated prolonged survival and displayed host-derived vascularization, thereby validating *in vivo* maturation. Collectively, HLCs provides a rapid and physiologically relevant liver model, with HLOs offering superior utility for disease modelling, therapeutic evaluation, and regenerative applications due to their enhanced functional and physiological relevance.

## 1 Introduction

Chronic liver diseases represent a major global health and economic burden, with metabolic dysfunction-associated steatotic liver disease (MASLD) and viral hepatitis infections as the predominant etiologies [1, 2]. As liver diseases continue to increase in prevalence, there is an urgent need for the development of effective tools to evaluate metabolism and toxicity during drug development [2, 3]. However, current *in vitro* and *in vivo* hepatic models, such as the human hepatoma cell lines (HepG2, Huh-7, and HepaRG) and animal models, fail to fully replicate human liver physiology because of chromosomal abnormalities [4], functional immaturity [5], and interspecies differences [6, 7]. Although the gold standard for *in vitro* hepatic models, primary human hepatocytes (PHHs), have overcome the limitations associated with supply and functional maintenance [8–10], their application to precision medicine and drug testing in various genetic backgrounds remains limited [11].

Recent advancement in human pluripotent stem cells (hPSCs) offers an opportunity to generate patient-specific hepatocytes like cells (HLCs) using two-dimensional (2D) cell-based [12] and 3D hepatic liver organoids (HLOs) *in vitro* platforms to study liver disease modelling [13–15]. The shortcomings of HLCs-2D overcome by HLOs-3D cultures that better resemble the intricate architectural and functional properties of *in vivo* tissues, such as cell density, presence of non-parenchymal cells, organization, and communication. Several studies have reported methods for differentiating hPSCs into hepatocyte-like cells (HLCs) and hepatic liver organoids (HLOs) [16–25]. Subsequently, these protocols are dependent on various cytokines (FGF2, BMP4, HGF, and KGF)[26, 27] and small molecules (CHIR99021and Dihexa) [28, 29] resulting in complex, labor intensive, cost-prohibitive workflows that limit scalability. Moreover, most of the reported methods predominantly differentiate cells into hepatic epithelial cell types, lacking essential supporting components such as pro-fibrotic and/or inflammatory cell types, and thus have limited capacity to model inflammatory disease [15, 30, 31]. These limitations led us to develop a streamlined 10-day protocol that utilizes substrate-directed morphogenesis: seeding hPSCs onto conventional tissue culture plates yields monolayer HLCs, while Elplasia™ 3D matrices promote self-organizing organoids in an extracellular matrix (ECM)-independent manner. The resulting dual-output platform generates scalable HLCs and HLOs exhibiting multicellular hepatic complexity along with functional hepatocytes. Both models demonstrate mature hepatic physiology, evidenced by sustained albumin secretion, CYP3A4 metabolic activity, glycogen storage, and free fatty acid (FFA) uptake. However, HLOs best recapitulate the cellular complexity and microenvironmental context of the native liver. Further, subcapsular transplantation of HLOs in NOD-SCID mice, with sustained expression of ALB and HNF4A, validates their physiological relevance for disease modelling and therapeutic screening. Collectively, this dual-output platform represents a reproducible, scalable, and clinically relevant *in vitro* system that bridges the gap between reductionist 2D models and the multicellular complexity of the native liver, offering a powerful tool for dissecting disease mechanisms, identifying therapeutic targets, and advancing personalized medicine for hepatic disorders.

## 2 Results

### 2.1 Establishment of a hPSC-derived HLCs

Induction of DE from hPSCs is a critical first step for subsequent efficient hepatic differentiation [29]. Prior to the initiation of differentiation (Day 0), the hPSCs were assessed for the expression of pluripotency markers (NANOG and OCT4) (Fig. 2B). For DE induction, the hPSCs were treated with CHIR99021, and by the end of the induction period (Day 2), the cells displayed definitive endoderm morphology and expressed the endodermal markers FOXA2 and SOX17, as confirmed by immunostaining and flow cytometry analysis (Fig. 2B,D). Around 90 ± 3.2 % of hPSCs-derived DE cells were positive for SOX17 marker, confirming efficient induction of DE cells (Fig. 2D). Quantitative gene expression analysis shows upregulation of both endodermal (FOXA2 and SOX17) and mesodermal markers (BRACHYURY, MESP1, TBX3) in DE, supporting successful differentiation towards the DE lineage (Fig. 2C). At the end of the homogeneous DE induction, the cells were treated with 1% DMSO for 5 days to specify a hepatic fate. The cells show a rapid change in morphology and a spurt of proliferation shortly after the treatment. On day 7, cells exhibited a typical hepatoblast (HE) morphology (Fig. 2A) and expressed HNF4A and AFP, as evidenced by immunostaining and quantitative gene expression analysis (Fig. 2B,E). While the exact role of DMSO in this protocol is unclear, it is critical.

**Fig. 1.**
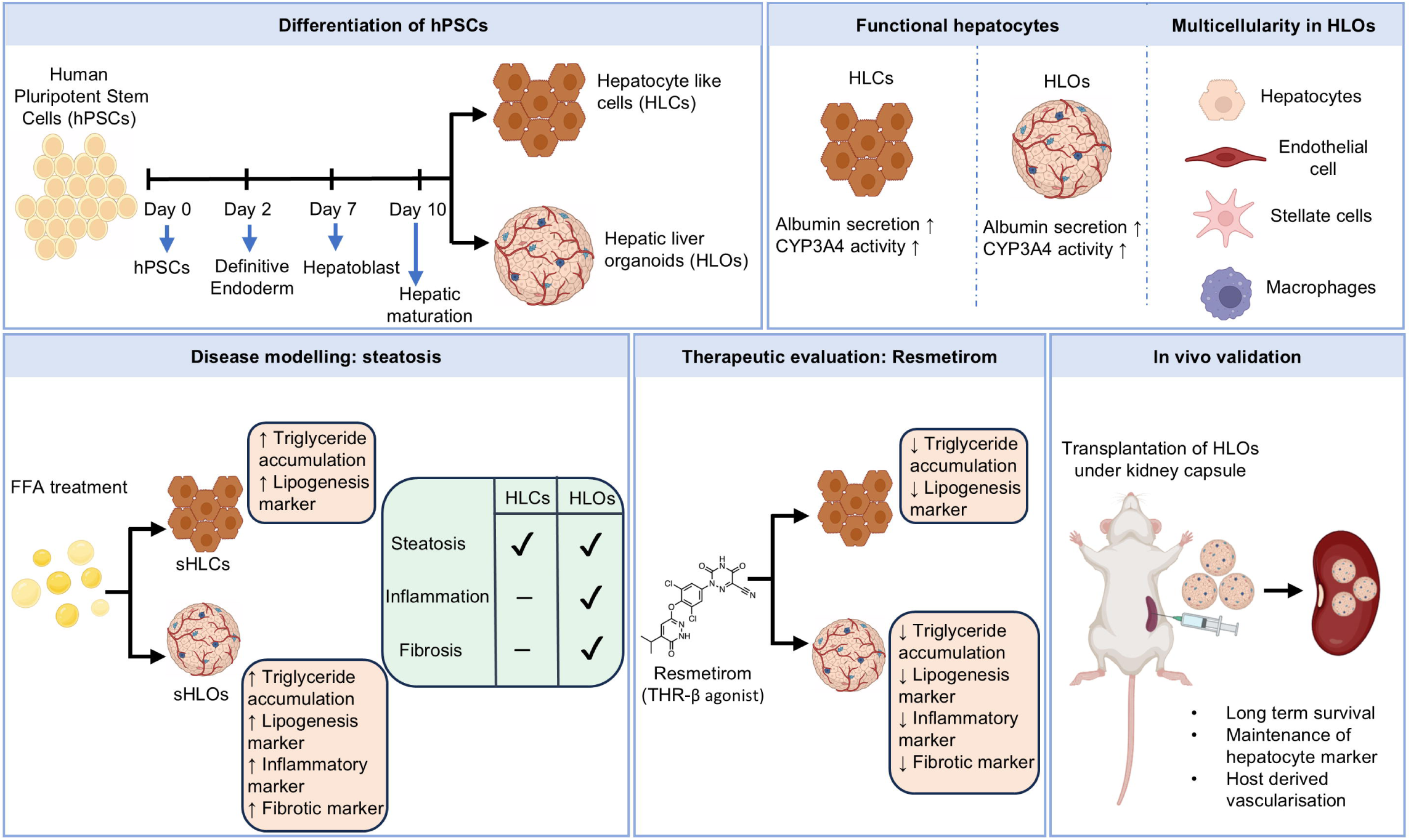
Schematic illustration of generation of hepatocyte-like cells (HLCs) and hepatic liver organoids (HLOs) from human pluripotent stem cells (hPSCs), and their subsequent used in metabolic dysfunction-associated steatotic liver disease (MASLD) modelling

**Fig. 2.**
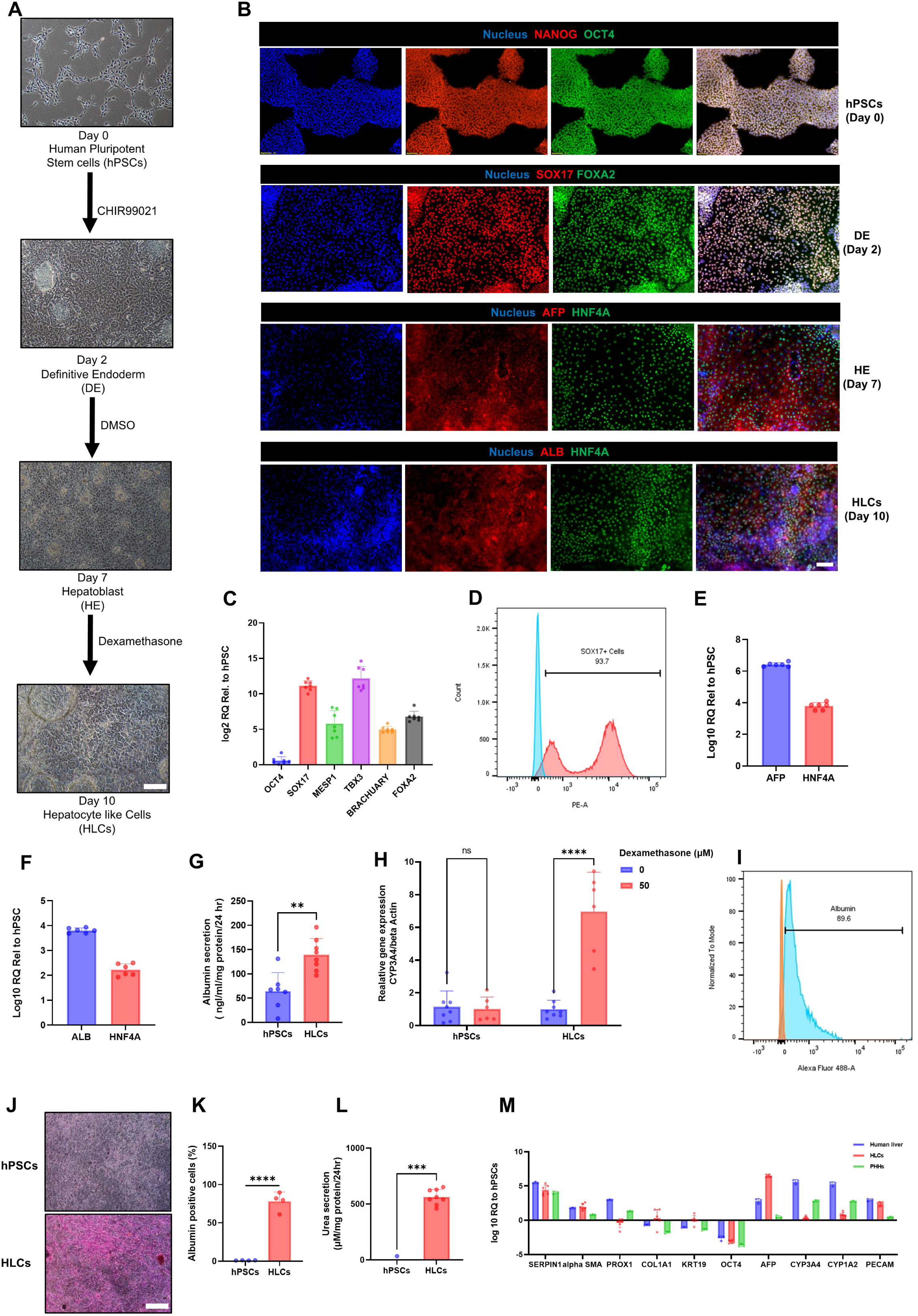
Characterization of hPSCs derived Hepatcyte like cells (HLCs). (A) Bright-field micrographs of directed differentiation of hPSCs into HLCs. Scale bar, 100 μm. (B) Immunostaining of hPSCs markers (OCT4, NANOG) on day 0, DE markers (FOXA2, SOX17) on day 2, HE markers (HNF4A, AFP) on day 7, HLCs markers (HNF4A, ALB) on day 10. Scale bar, 100 μm. C) Quantitative gene expression analysis of mesoderm and endoderm markers in day 2 cells. (D) Flow cytometry analysis of DE cells confirming SOX17-positive cells on day 2. (E) Quantitative gene expression analysis of HE markers on day 7. (F) Quantitative gene expression analysis of HLCs markers on day 10. (G) Analysis of albumin secretion by hPSCs and HLC. (H) Quantitative gene expression analysis of CYP3A4 upon induction with dexamethasone (CYP3A4 inducer) in hPSCs and HLCs. (I and K) Flow cytometry analysis confirming ALB-positive cells on day 10. (J) PAS staining demonstrating glycogen storage (dark pink) in hPSCs and HLCs. Scale bar, 100 μm. (L) Analysis of urea secretion hPSCs and HLCs. (M) Quantitative gene expression analysis of HLCs, PHH (primary human hepatocytes), and pooled human liver. Data represent mean ±SD. n.s., not significant, *p < 0.05, **p > 0.01, ***p > 0.001, ****p > 0.0001; Unpaired t-test (G, K, L), Two-way ANOVA (H).

At day 7 of differentiation, the cells were further treated for 3 days with dexamethasone, a glucocorticoid mimetic, for hepatocyte maturation. At the end of the maturation (day 10), hepatocyte-like cells (HLCs) displayed a typical cobblestone morphology resembling primary hepatocytes (Fig. 2A). These cells also demonstrated expression of hepatocyte markers ALB and HNF4A in immunostaining and quantitative gene expression analysis (Fig. 2B,F).

We then assessed hepatic functions of HLCs. Secretion of albumin (ALB) is a common indication of hepatic maturation. The HLCs secrete significant amounts of ALB as compared to hPSCs (Fig. 2G), confirming hepatic maturation. More importantly, we assessed drug metabolism by the CYP3A cytochromes, a major family of liver detoxifying enzymes in hepatocytes. To more specifically address the activity of the CYP3A4, a marker of mature hepatocytes, we incubated HLCs with a CYP3A4-inducer. The observed upregulation of the CYP3A4 gene after treatment with inducer confirms the activity of the CYP3A4 in HLCs (Fig. 2H). Additionally, urea secretion was measured and found significant amount of urea secretion by HLCs as compared to hPSCs (Fig. 2L). We found that more than 80% of the cells were positive for ALB, as determined by flow cytometry analysis, indicating a high proportion of successful hPSCs differentiation into HLCs (Fig. 2I,K). We also observed glycogen storage in HLCs through periodic acid-schiff (PAS) staining (Fig. 2J). Next, gene expression analysis between HLCs, human liver tissue, and primary human hepatocytes (PHHs) revealed difference in AFP and ALB gene (Fig. 2M). Notably, HLCs exhibited elevated AFP expression compared to PHHs and human liver RNA, which is typically associated with fetal or immature hepatocytes rather than fully mature adult hepatocytes. Conversely, ALB gene expression in HLCs was lower than that observed in PHHs and human liver RNA. Interestingly, we found expression of endothelial cell marker (PECAM) and hepatic stellate cell marker (alpha SMA) (Fig. 2M). It suggests the presence of both parenchymal (hepatocyte cells) and non-parenchymal (hepatic stellate cells, endothelial cells) cells at day 10 of differentiation stage.

### 2.2 Characterisation of diverse cell population

Our findings indicated that approximately 80% of the cells expressed hepatocyte marker ALB on day 10. To further characterize the remaining ∼20% of cells, we investigated the presence of non-parenchymal cell populations, evidenced by the observed expression of both parenchymal and non-parenchymal gene markers. We performed a quantitative fluorescent antibody-based profiling by flow cytometry with epithelial marker EpCAM, the stellate cell marker CD166/ALCAM, and the Kupffer cell markers CD68 and endothelial marker CD31. The frequency of EpCAM+ cells was 51 ± 0.88% of total cells, whereas EpCAM− CD166+ and EpCAM− CD31+ expressing cells was 22.4 ± 1.53%, and 3.36 ± 0.46% in EpCAM− cells, respectively (Fig. 3B). However, no population of EpCAM−CD68+ was observed, indicates absence of kupffer cells. These findings were further supported by immunostaining results (Fig. 3A). This protocol, which starts with the differentiation of DE into HLCs, could result in reproducible differentiation and reduced heterogeneity of HLCs and non-parenchymal cells.

**Fig. 3.**
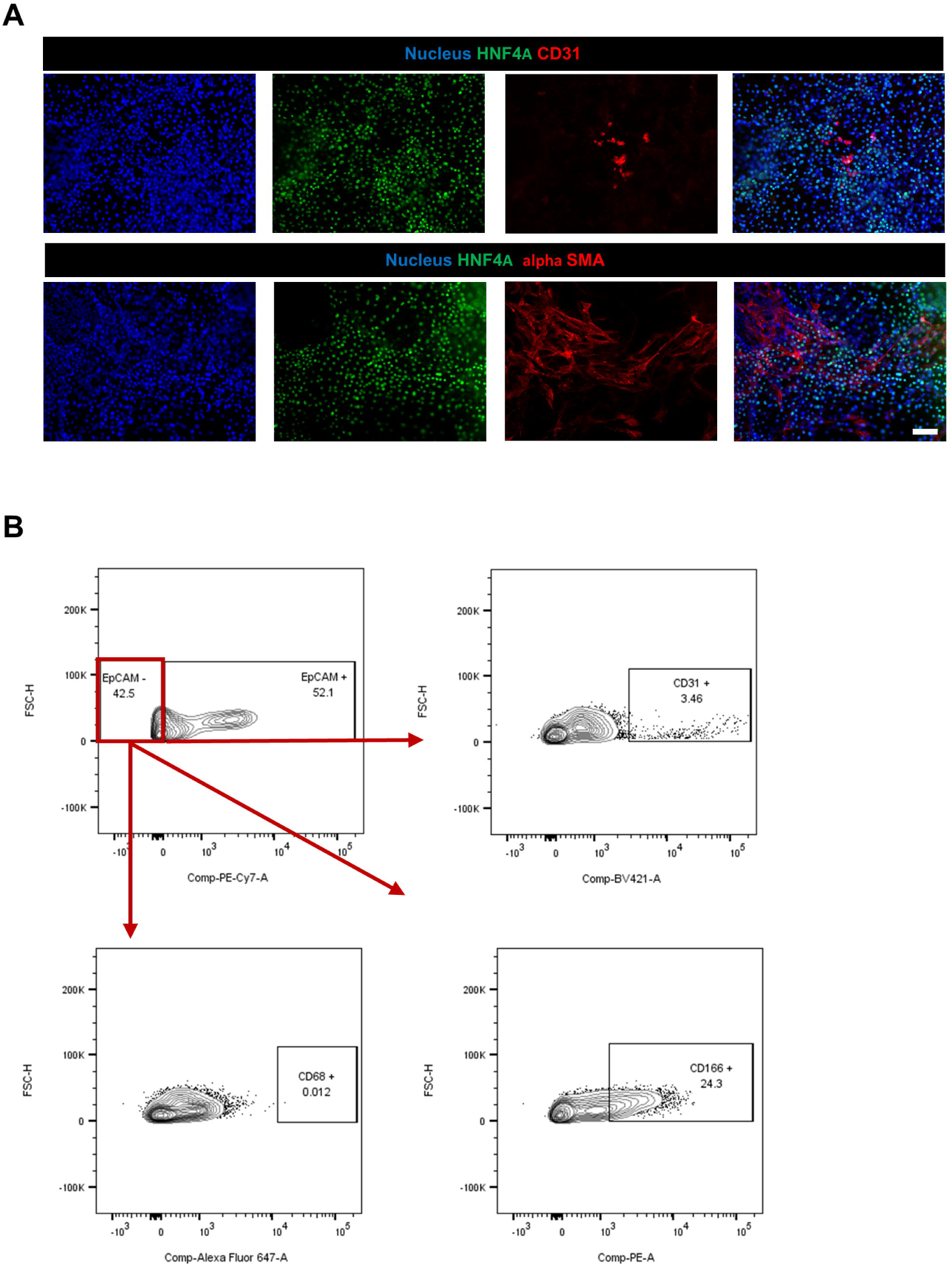
Characterization of non-parenchymal cells of liver on day 10. (A) Immunostaining of hepatic stellate (alpha SMA) and endothelial cells (CD31) on day 10. Scale bar, 100 μm. (B) Flow cytometry analysis of EpCAM-/CD166+ (stellate cells) and EpCAM-/CD31+ (endothelial cells) cells on day 10.

### 2.3 HLCs can be induced to steatotic HLCs for MASLD modelling

To better simulate the MASLD condition, we established an *in vitro* model that promotes steatosis in HLCs and defined FFA treated HLCs as steatotic HLCs (sHLCs). It involved treating the cells with a 2:1 ratio of oleic acid (OA) and palmitic acid (PA) along with vehicle. First, we examined the effect of FFA on the cell viability of HLCs. Interestingly, we observed no cytotoxicity in 0.2, 0.4, 0.8, 1.6 and 3.2 mM concentration of FFA for 48 h (Fig. 4B). Next, we observed a significant dose dependent increase in lipid accumulation compared to the vehicle, as visualised by BODIPY and quantified by Oil red O (ORO) staining and (Fig. 4A,C) in sHLCs. Next, we quantified the neutral lipid (Nile Red) and ROS (2’,7’-dichlorodihydrofluorescein diacetate (H2DCFDA) by flow cytometry in sHLCs and found a significant dose dependent increase in neutral lipids and ROS as compared to the vehicle (Fig. 4E-H). Further, intracellular triglycerides (TGs) levels were quantified in the sHLCs and found dose dependent increased in sHLCs as compared to vehicle (Fig. 4D).

**Fig. 4.**
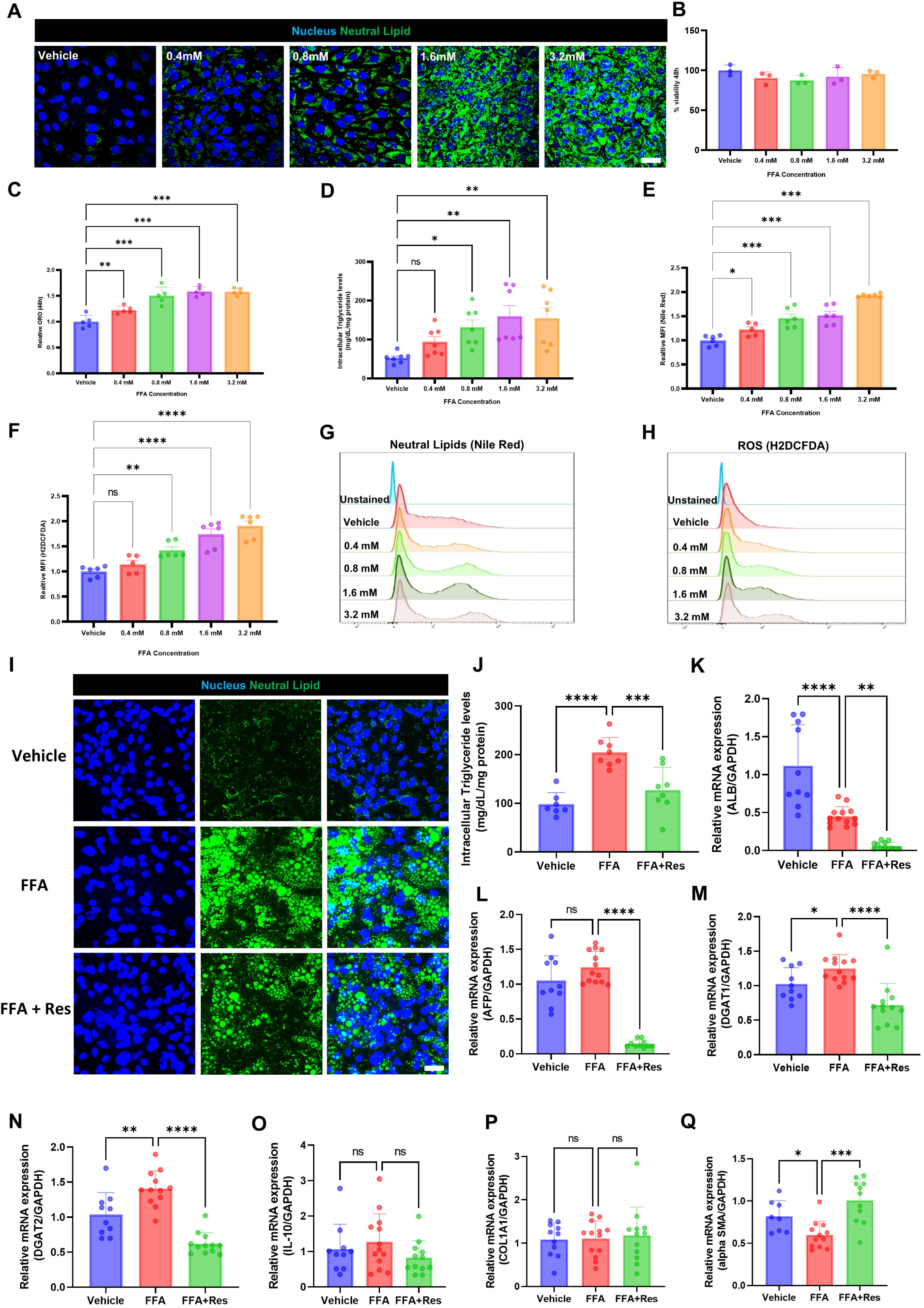
Modelling MASLD phenotype using HLCs. (A) Confocal micrographs of HLCs treated with increasing concentrations of FFA (0.2–3.2 mM), stained for neutral lipids (BODIPY, green) and nuclei (Hoechst 33342, blue). Scale bar = 20 μm. (B) Cell viability analysis of sHLCs following FFA treatment. C) Relative quantification of Oil Red O staining in FFA-treated HLCs. (D) Quantification of triacylglycerol (TGs) synthesis and storage in FFA-induced HLCs after 48 h of treatment. (E, G) Flow cytometric analysis of neutral lipid accumulation in sHLCs quantified as mean fluorescence intensity (MFI) following Nile Red staining. (F, H) Flow cytometry analysis of ROS quantified by H2DCFDA staining of sHLCs. (I) Confocal micrographs of stained for neutral lipids (BODIPY, green) and nuclei (Hoechst 33342, blue), Scale bar = 20 μm and (J) quantification of TGs in HLCs treated with FFAs for 120 h, followed by resmetirom treatment for an additional 120 h. (K-L) Quantitative gene expression analysis of hepatocyte-specific markers; (M–N) lipid metabolism-associated genes; (O) inflammatory response markers; and (P–Q) fibrotic markers. Data represent mean ±SD. n.s., not significant, *p < 0.05, **p > 0.01, ***p > 0.001, ****p > 0.0001; one-way ANOVA test (B-F, J-Q). FFA- Free fatty acids.

Resmetirom, a selective agonist of thyroid hormone receptor beta (THRβ), has received approval for the treatment of metabolic dysfunction-associated steatohepatitis (MASH) and moderate to advanced hepatic fibrosis [32] [33]. To evaluate the potential of the developed sHLCs model for drug screening, we established a curative model of sHLCs using Resmetirom. To induce a curative model, HLCs were treated with 0.8 mM FFA for 120 h, followed by treatment with resmetirom at concentrations of 100 µM in the presence of FFA for an additional 120 h [34]. Treatment with resmetirom led to a reduction in TGs levels (Fig. 4J), accompanied by decreased neutral lipid accumulation as evidenced by reduced BODIPY staining (Fig. 4I). Moreover, treatment with FFA resulted in an increased expression of the lipogenesis markers DGAT1 and DGAT2, which was subsequently reduced following resmetirom treatment (Fig. 4M,N). Interestingly, resmetirom treatment resulted in a reduction in the expression of hepatic markers ALB and AFP in HLCs compared to vehicle suggesting a shift in hepatocyte function (Fig. 4K,L). Furthermore, there was no significant induction of IL-10, COL1A1, and alpha SMA upon FFA treatment, suggesting a lack of inflammatory and fibrotic responses (Fig. 4O-Q). These findings suggested that developed hPSCs derived HLCs could be effectively used as a model for MASLD.

### 2.4 Metabolomics and associated pathway analysis of sHLCs

We performed an untargeted metabolomic analysis of the sHLCs to examine changes in metabolites associated with MASLD. Using an untargeted metabolomics profiling approach, 452 metabolites were identified. The PCA plot showed a clear separation between the vehicle (BSA) and sHLCs (Fig. 5A). The volcano plot highlights differential metabolite species between BSA and sHLCs (Fig. 5B). Notably, sHLCs exhibited significant changes in lipid-associated metabolites, including increased levels of lysophosphatidylethanolamine (LPE 18:2) and altered inositol metabolism (Fig. 5C,D). In addition, multiple lipid species, including phosphatidylcholine (PC 18:1), phosphatidylinositol (PI 36:2), and fatty acids such as oleic acid (FA 18:1) and eicosenoic acid (FA 20:1), were significantly increased as compared to BSA, reflecting enhanced lipid accumulation (Fig. 5F–J). Conversely, a marked reduction in glutathione and phosphocreatine levels was observed, indicating impaired redox balance and disrupted cellular energy homeostasis (Fig. 5E,K). The decrease in creatine further supports dysregulation of the phosphocreatine energy buffering system in sHLCs [35]. Collectively, these metabolic alterations recapitulate key features of MASLD and highlight potential metabolic signatures associated with disease progression.

**Fig. 5.**
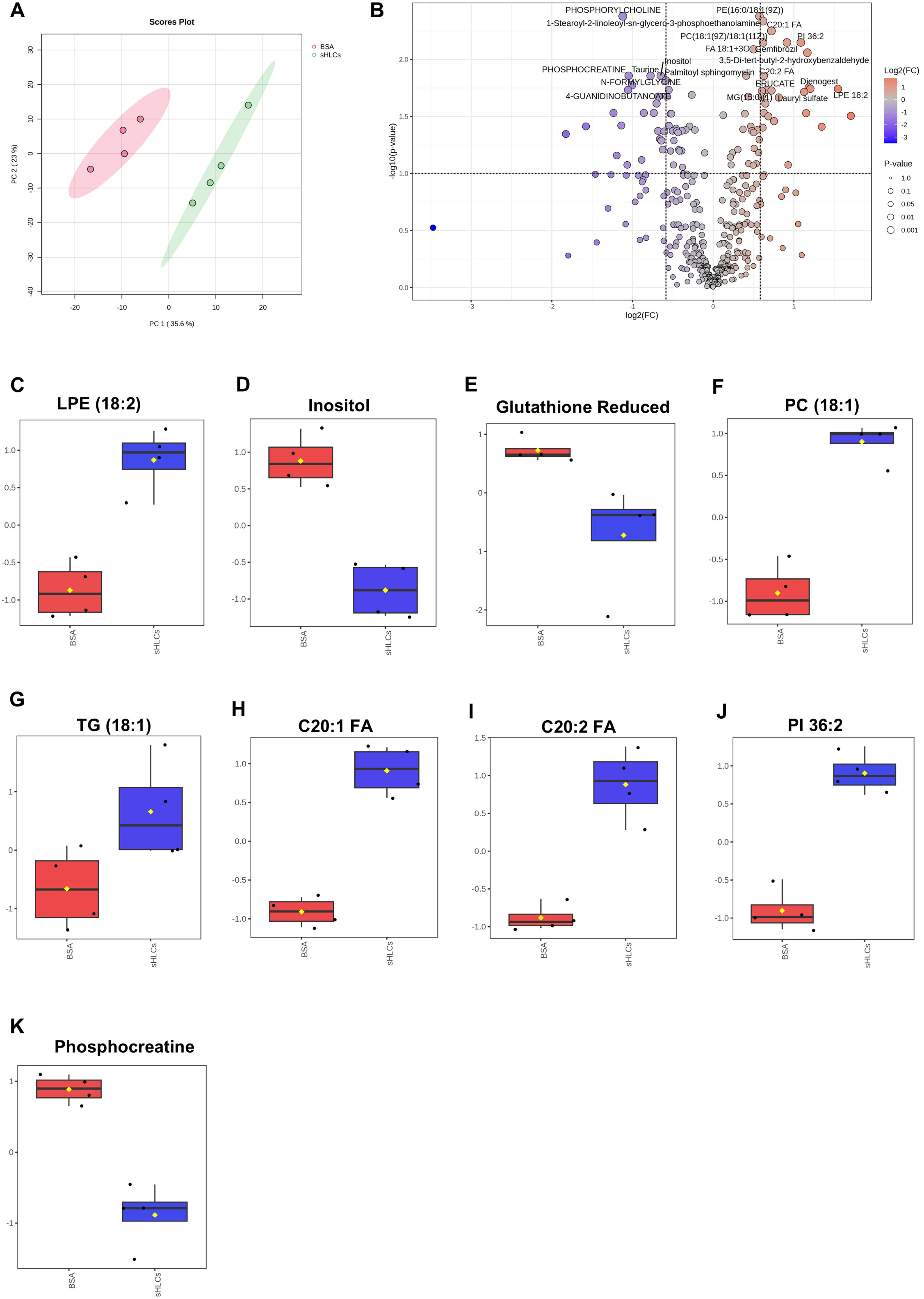
Metabolite profiling of sHLCs. Two-dimensional score plot of principal component analysis (PCA) comparing control HLCs and sHLCs. (B) Volcano plot showing differential metabolite expression between control and FFA-treated HLCs. (C–L) Box plot-based quantification of selected metabolites and lipid species in control HLCs and sHLCs, including (C) lysophosphatidylethanolamine (LPE), (D) phosphatidylinositol (PI), (E) glutathione, (F) phosphatidylcholine (PC), (G) triglycerides (TG), (H–I) fatty acids (FAs), (J) phosphatidylinositol (PI) species and (K) phosphocreatine. The middle line in the box equals the median; the top and bottom lines are the first and third quartiles.

### 2.5 Establishment of a hPSC-derived HLOs

To evaluate whether the optimized 10-day differentiation protocol could be utilized for HLO generation, the protocol was applied under 3D conditions. Upon single-cell seeding in ultra-low attachment conditions, hPSCs rapidly self-aggregated to form compact spheroids within 24 h (Fig. 6A). These aggregates maintained the expression of pluripotency markers OCT4 and NANOG, indicating the preservation of pluripotency during early aggregation (Fig. 6B). Initially, the hPSCs (day 0) were differentiated into foregut spheroids through definitive endoderm specification using CHIR99021[36] [37]. This transition (day 2) resulted in the expression of both endoderm (FOXA2/SOX17) and mesoendoderm (BRACHYURY) markers, as evaluated by immunostaining and quantitative gene expression analysis (Fig. 6B,C). On day 2, cells were treated with differentiation media with 1% DMSO for 5 days, followed by differentiation media with dexamethasone treatment for 3 days to induce the hepatocyte differentiation process, thereby establishing human liver organoids (HLOs), on day 10. The HLOs exhibited features of hepatocyte-like maturation, including increased expression of ALB and HNF4A evaluated by both quantitative gene expression analysis and immunostaining (Fig. 6B,D). Functional characterization of HLOs demonstrated acquisition of key hepatic function, including a significant increase in albumin secretion as compared to hPSCs (Fig. 6E). Furthermore, treatment with a CYP3A4 inducer resulted in significant induction of CYP3A4 activity in HLOs, confirming the presence of CYP450 activity (Fig. 6F). Further quantitative gene expression analysis revealed stage-specific gene expression across day 7 and day 10 organoids compared to human liver RNA. The day 7 organoids exhibited expression of progenitor-associated markers, including AFP, PROX1, and KRT19, along with detectable COL1A1 and alpha SMA, indicating a heterogeneous and early hepatic state (Fig. 6D). By day 10, organoids showed increased expression of mature hepatic markers SERPINA1, CYP1A2, and CYP3A4, accompanied by reduced OCT4 expression, confirming hepatic differentiation (Fig. 6D). The PECAM expression suggested the presence of endothelial-like populations at day 10 (Fig. 6D).

**Fig. 6.**
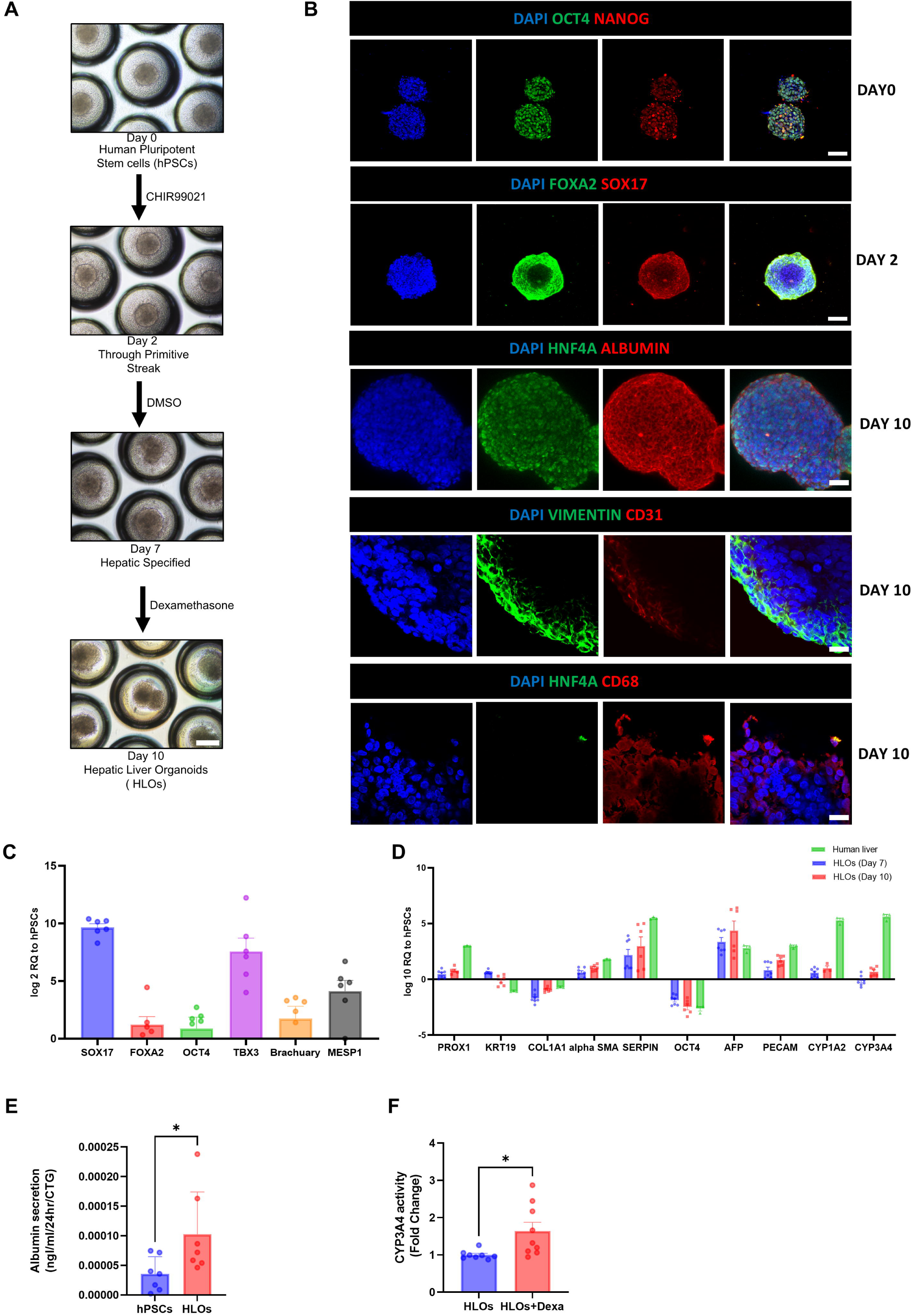
Characterization of hPSC-derived hepatic liver organoids (HLOs). (A) Bright-field micrographs showing the directed differentiation of human pluripotent stem cells (hPSCs) into hepatic liver organoids (HLOs). Scale bar = 100 μm. (B) Immunostaining of pluripotency markers (OCT4 and NANOG) on day 0; DE markers (FOXA2 and SOX17) on day 2; and hepatic and non-parenchymal cell markers on day 10, including hepatocyte markers (HNF4A and ALB), stellate cell marker (Vimentin), endothelial cell marker (CD31), and Kupffer cell marker (CD68). Scale bars = 100 μm unless otherwise indicated; CD31 and CD68, 20 μm. (C) Quantitative gene expression analysis of mesodermal and endodermal markers on day 2. (D) Quantitative gene expression analysis of HLOs at day 7 and 10 compared with pooled human liver samples. (E) Quantification of albumin secretion by HLOs. (F) CYP3A4 enzymatic activity in HLOs following induction with dexamethasone, a CYP3A4 inducer. Data represent mean ±SD. n.s., not significant, *p < 0.05, **p > 0.01,***p > 0.001, ****p > 0.0001; Unpaired t-test (E and F).

### 2.6 Characterisation of diverse cell populations and *in vivo* maturation of HLOs

We performed a quantitative fluorescent antibody-based profiling of the HLOs by flow cytometry analysis with epithelial marker EpCAM, the stellate cell marker CD166/ALCAM, and the Kupffer cell markers CD68 and endothelial marker CD31. The frequency of EpCAM+ cells was 35.20 ± 0.45% of total cells, whereas EpCAM−CD166+, EpCAM−CD68+, and EpCAM−CD31+ expressing cells was 65.58 ± 1.12%, 13.89 ± 0.28%, and 10.84 ± 0.39% in EpCAM− cells, respectively (Fig. 7A). Further immunostaining analysis of the HLOs demonstrated that most cells express the hepatocyte markers ALB and HNF4A. Additionally, the stellate cell marker vimentin was observed in HLOs with a spindle-shaped morphology, while the endothelial (CD31) and Kupffer (CD68) markers were detected at frequencies comparable to those quantified by flow cytometry analysis (Fig. 6B).

**Fig. 7.**
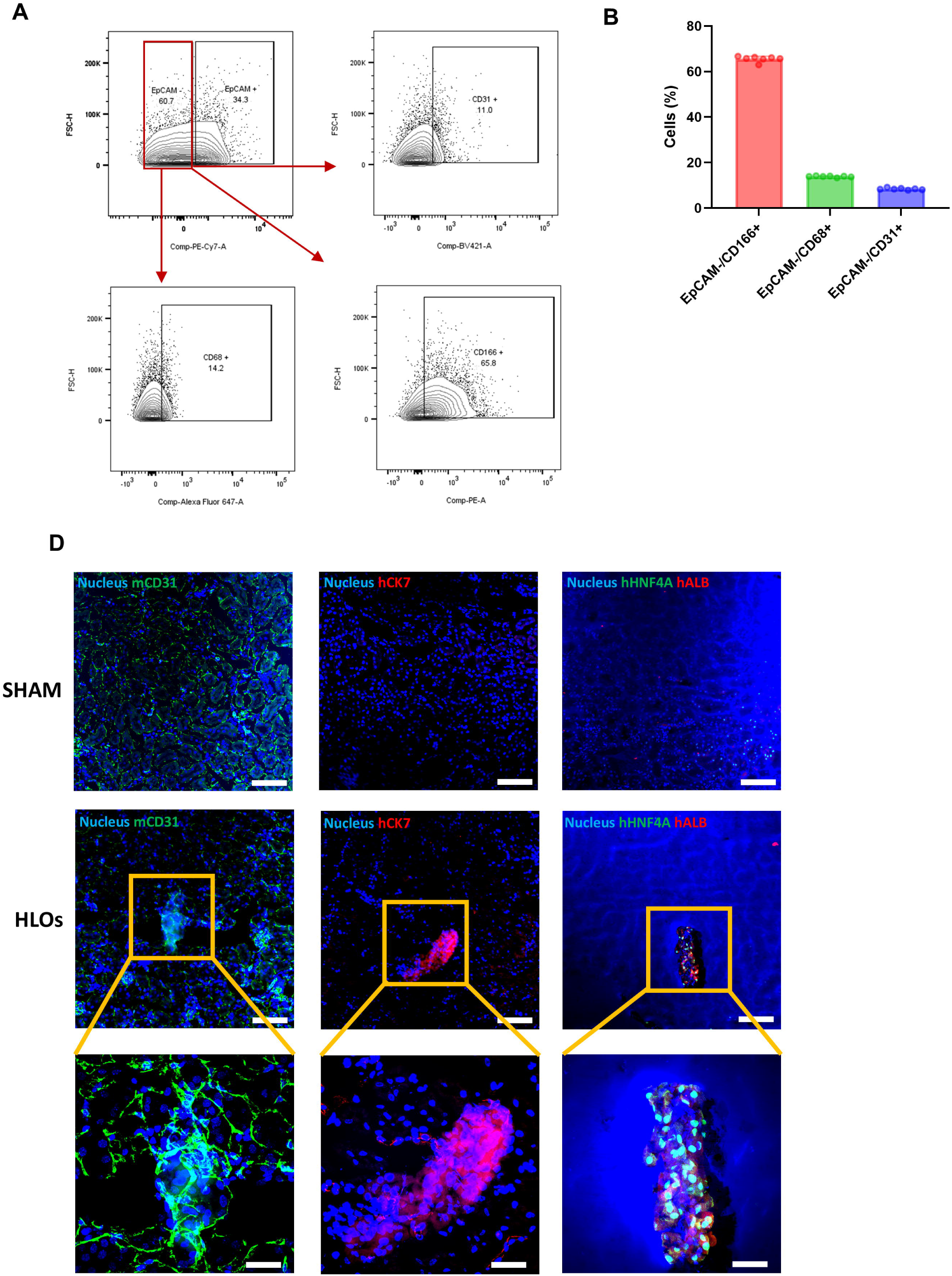
Characterization of non-parenchymal cell populations in HLOs. (A–B) Flow cytometric analysis of non-parenchymal cell populations in hepatic liver organoids (HLOs), including EpCAM-/CD166+ stellate cells, EpCAM-/CD31+ endothelial cells, and EpCAM-/CD68+ Kupffer cells. (C) Immunostaining of cryosections from mouse kidney HLO transplants, showing human hepatic cell populations (hCK7, hHNF4A, and hALB) and mouse endothelial cells (mCD31). Scale bar = 100 μm for full images and 20 μm for enlarged inset images (bottom) showing detailed cellular architecture and cell-cell interactions.

To evaluate the *in vivo* maturation of HLOs, these organoids were transplanted beneath the kidney capsule of immunodeficient NOD-SCID mice. Subsequent immunostaining of the explanted grafts revealed a strong expression of hepatocyte markers ALB and HNF4A in transplanted HLOs at week 4 (Fig. 7D), thereby confirming the preservation of hepatocyte identity in *in vivo*. Additionally, the presence of the biliary-associated marker CK7 was detected, indicating the maintenance of hepatic epithelial heterogeneity (Fig. 7D). Notably, the detection of host-specific (mouse) CD31 within the transplanted HLOs indicates host-derived endothelial cell infiltration and subsequent vascularization (Fig. 7D), thereby supporting the growth and survival of HLOs within the *in vivo* microenvironment.

### 2.7 HLOs can be induced to steatotic HLOs for MASLD modelling

To establish an *in vitro* model of MASLD, HLOs were treated with FFA (800µM), leading to significant intracellular lipid accumulation as compared to vehicle, as verified by BODIPY staining and TGs level (Fig. 8A,G) and we defined FFA treated HLOs as steatotic HLOs (sHLOs). These lipid droplets appeared enlarged and densely distributed in sHLOs as compared to vehicle, indicating excessive TGs storage and a steatotic phenotype. The resmetirom treatment markedly attenuated lipid accumulation, as evidenced by reduced BODIPY staining and TGs level. (Fig. 8A,G). This lipid accumulation was linked to altered transcriptional alterations related to MASLD progression. Specifically, quantitative gene expression analysis showed an increase in the expression of lipogenesis marker (DGAT1 and DGAT2), fibrosis markers (alpha SMA, COL1A1) and inflammatory marker (IL-10) (Fig.8 B-F). Notably, resmetirom treatment to sHLOs resulted in decreased expression of DGAT1, DGAT2, COL1A1, and alpha SMA, while also restoring IL-10 levels as compared to sHLOs (Fig. 8B-H). This suggests a partial reversal of both fibrotic and inflammatory responses, highlighting the potential of sHLOs as a physiologically relevant platform for modelling steatohepatitis and evaluating therapeutic interventions.

**Fig. 8.**
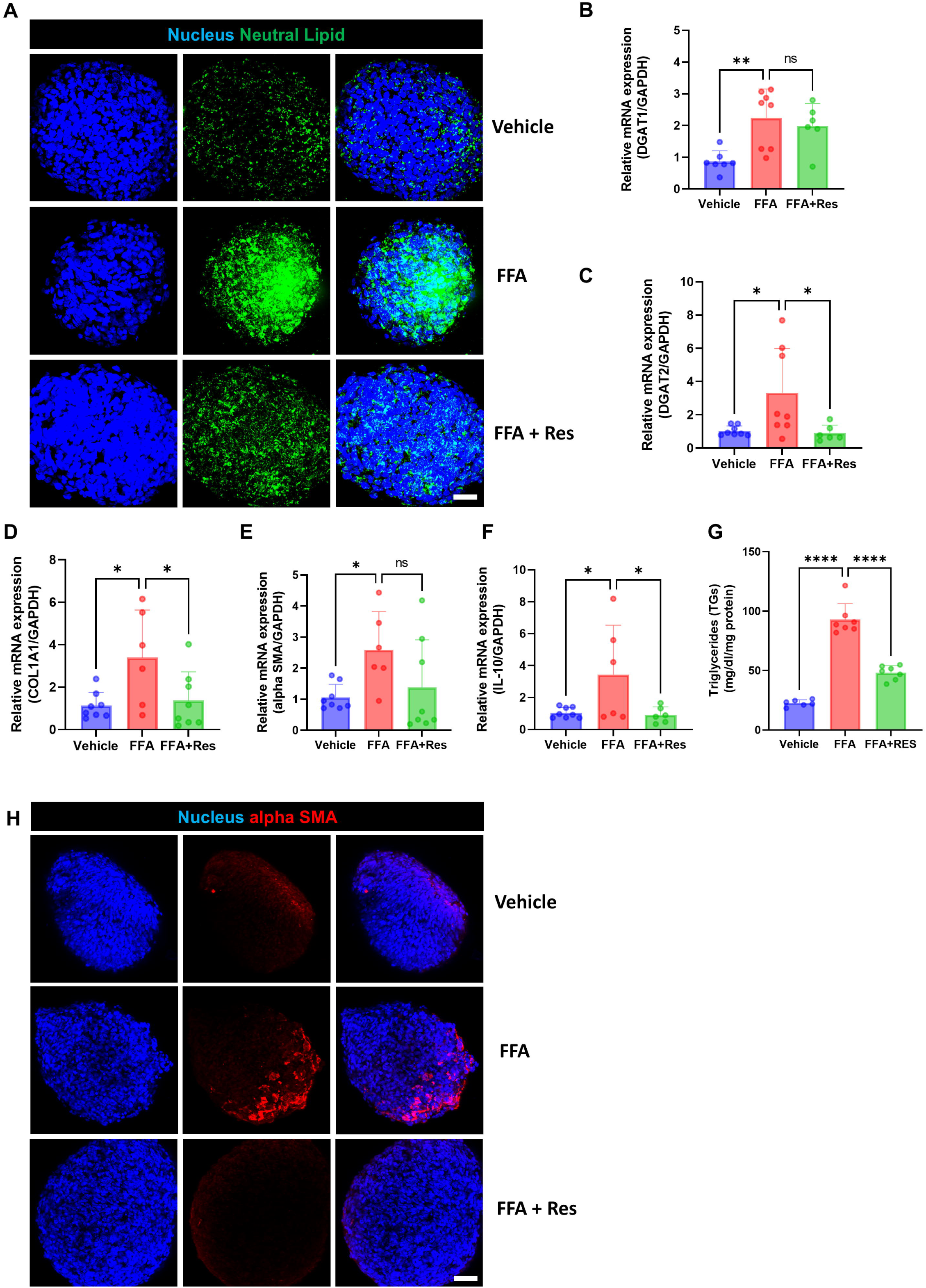
Modelling MASLD phenotype using HLOs. (A) Confocal micrographs showing lipid accumulation and (G) quantification of TGs in HLOs treated with FFA (FFA) for 120 h, followed by resmetirom treatment (FFA+Res) for an additional 120 h. Scale bar = 20 μm. Quantitative gene expression of (B-C) lipid metabolism-associated genes, (D) Inflammatory response markers, and (E-F) fibrotic markers. (H) Immunostaining of alpha SMA in FFA and FFA+Res. Scale bar = 100 μm. Data represent mean ± SD, n.s., not significant, *p < 0.05, **p > 0.01, ***p > 0.001, ****p > 0.0001; one-way ANOVA test (B-F). FFA- Free fatty acid, Res- Resmetirom.

### 2.8 Comparative analysis of 2D HLCs and 3D hepatic organoids

A comparative analysis of HLCs and HLOs both derived using an identical differentiation protocol, revealed significant differences in cellular organization and functional outcomes as described in Table 1.

**Table 1.** Comparative analysis of 2D hepatocyte-like cells (HLCs), 3D hepatic liver organoids (HLOs).

| Feature | 2D HLCs | 3D HLOs |
| --- | --- | --- |
| **1. Differentiation protocol** | 10-day small molecule–driven protocol | 10-day small molecule–driven protocol |
| a. **Hepatic markers** (ALB, HNF4A, CYP3A4) | Expressed | Expressed |
| **b. Cellular composition** | Predominantly hepatocyte-like cells, hepatic stellate cells, and endothelial cells | Multicellular (hepatocyte, hepatic stellate cells, kupffer cells, and endothelial cells) |
| **c. Structural organization** | Monolayer, limited architecture | 3D architecture with cell–cell interactions |
| **d. Functional maturation**<br>(albumin secretion, CYP activity) | Present | Present |
| **e. *In vivo* relevance** | Limited | High (supports transplantation studies) |
| **2. MASLD modelling** | Lipid accumulation observed | Lipid accumulation observed along with steatohepatitis phenotype |
| **a. Fibrotic response** (alpha SMA, COL1A1) | Absent | Strong induction |
| **b. Inflammatory response** (IL-10) | Absent | Regulated response |
| **c. Drug response** (Resmetirom) | Observed therapeutic response | Enhanced therapeutic response |
| **d. Overall suitability** | Scalable, drug screening | Advanced liver model, physiologically relevant disease modelling and drug screening |

## 3 Discussion

Here, we present a rapid, small molecule–driven platform for generating hepatocyte-like cells (HLCs) and hepatic liver organoids (HLOs) from hPSCs that recapitulate key features of MASLD. The system achieves efficient hepatic differentiation within 10 days and supports multicellular organization in 3D, including hepatocyte and non-parenchymal populations in HLOs compared to HLCs. Upon FFA challenge, both models develop steatosis, while HLOs uniquely capture advanced disease phenotypes, including fibrotic and inflammatory responses (steatohepatitis) (Fig. 1). Functional and transcriptional analyses confirm hepatic maturation and metabolic competence, while therapeutic intervention with resmetirom attenuates lipid accumulation and partially reduces disease-associated signatures.

Efficient CHIR99021-mediated induction resulted in near-homogeneous definitive endoderm, consistent with Wnt-driven lineage commitment essential for hepatic differentiation [27, 38]. Subsequent DMSO-driven specification and dexamethasone-mediated maturation exhibiting key hepatic features, including albumin secretion, urea synthesis, glycogen storage, and inducible CYP3A4 activity, as described in prior reports [8]. The transient upregulation of mesodermal genes suggests progression through a mesendodermal intermediate, which may facilitate lineage plasticity and enhance differentiation efficiency [24].

The liver, composed of hepatocytes for nutrient metabolism, kupffer cells as primary resident macrophages, hepatic stellate cells for regulating liver fibrosis, and liver sinusoidal endothelial cells for facilitating nutrient exchange, is further structured by cholangiocytes lining bile ducts for bile production [39]. We identified cell population of stellate and endothelial cells along with hepatocytes upon differentiation. The embryogenic origin of stellate cells has been unclear, with evidence suggesting they originate from either the endoderm or the septum transversum, which forms from cardiac mesenchyme during the invagination of the hepatic bud [40]. Interestingly, hPSC induction with CHIR99021 resulted in the co-expression of mesoderm and endoderm markers at the DE stage, potentially explaining the generation of mesoderm-derived stellate and endothelial cells alongside endoderm-derived hepatocytes in this protocol [24]. This coexistence of parenchymal and non-parenchymal populations may enhance physiological relevance.

The establishment of a steatotic HLC (sHLC) model effectively recapitulates key early features of MASLD, including lipid accumulation, oxidative stress, and TG overload [41]. The absence of cytotoxicity across a broad FFA concentration range indicates that the observed phenotypes are driven by metabolic dysregulation rather than cell death, consistent with early-stage steatosis. The dose-dependent increase in neutral lipids, ROS, and intracellular TG reflects an imbalance between lipid accumulation and clearance, a central mechanism underlying MASLD progression [42] [43] [44]. Therapeutic intervention with Resmetirom significantly reduced lipid accumulation and downregulated lipogenic genes (DGAT1, DGAT2), confirming restoration of metabolic homeostasis. Interestingly, the reduction in ALB and AFP expression suggests a shift in hepatocyte functional state, potentially due to metabolic reprogramming rather than loss of viability. Notably, the absence of significant induction of inflammatory (IL-10) and fibrotic (COL1A1, alpha SMA) markers indicates that this sHLCs model primarily captures early stages of steatosis without progression to advanced disease phenotypes. Collectively, these findings validate the sHLC system as a robust and scalable platform for modelling early MASLD and evaluating metabolic disorders therapeutics.

Untargeted metabolomic profiling revealed significant alterations in lipid species, including increased LPE (18:2), phosphatidylcholine, phosphatidylinositol, and fatty acids, consistent with lipid dysregulation in MASLD. Earlier studies demonstrated that elevated levels of phosphatidylethanolamines (PEs) increase susceptibility to the disease progression of obesity associated with MASLD, likely through a causal cascade impact on liver function [45] [46]. Inositols (INS) are ubiquitous polyols involved in numerous physiological functions. Alterations in INS metabolism appear to play a role in diseases involving insulin resistance, such as diabetes and polycystic ovary syndrome, and INS deficiency is associated with increased fatty liver in animals [47]. High amounts of free cholesterol in the liver reduce the activity of the 2-oxoglutarate carrier. As a result, less GSH is translocated from the cytosol into mitochondria, favouring ROS accumulation and the consequent lipid peroxidation. The mitochondrial GSH transporter 2-oxoglutarate carrier is involved in MASH development [48]. A close relationship exists between liver diseases and GSH, which supports our finding in metabolite profiles in the sHLCs model.

Implementation of the 10-day protocol in a 3D culture system enabled the rapid generation of hPSC-derived HLOs, characterized by efficient endoderm specification followed by robust hepatic differentiation. By day 10, HLOs exhibited increased expression of ALB and HNF4A, along with functional maturation indicated by albumin secretion and inducible CYP3A4 activity. The gene expression profiling showed a transition from progenitor to mature hepatic signatures, closely resembling human liver, albeit with residual fetal traits [37, 49]. The cellular profiling confirmed the presence of hepatocyte, stellate, endothelial, and minimal Kupffer-like populations, indicating multicellular hepatic complexity. Moreover, the HLOs could be successfully transplanted into mice and engrafted HLOs showed mice CD31-positive endothelial structures throughout large areas of the transplanted organoid, extending into the kidney parenchyma of the mouse. This process could only occur through vascularization from the organoids into the surrounding mouse tissue, supporting our claims of de novo vascularization [20, 24]. Collectively, these findings highlight enhanced maturation and cellular diversity, supporting their physiological relevance for the human liver microenvironment.

FFA treatment of HLOs induced lipid accumulation, as evidenced by enlarged lipid droplets establishing a steatotic phenotype (sHLOs) and also accompanied by upregulation of DGAT1 and DGAT2, indicating enhanced TGs synthesis. Increased alpha SMA expression suggested early fibrotic activation, while altered IL-10 levels reflected a compensatory inflammatory response similar to MASH or steatohepatitis phenotype [25, 50]. Collectively, these findings demonstrate that lipid overload drives both metabolic and fibrotic remodelling in HLOs. Notably, resmetirom treatment reduced lipid accumulation (TGs) and normalized gene expression, indicating partial reversal of MASLD-associated phenotypes.

Collectively, 3D-HLOs exhibited enhanced hepatic cellular complexity and tissue-like organization compared to 2D-HLCs, despite being generated using the same protocol. The current understanding is that dietary fat, FFAs from adipose tissue, and de novo liver lipogenesis triggered by carbohydrates are the main contributors to the pathophysiology of MASLD. These factors might further contribute to the progression of fibrosis by inducing insulin resistance and inflammation. Our system demonstrates that FFA alone are sufficient to realistically mimic MASLD *in vitro*. In future studies, our system could be used to investigate the precise role of other dietary and environmental factors in the pathogenesis of MASLD. The human-relevant platform, based on hPSC-derived hepatocytes/hepatic liver organoids (HLOs), provides a robust system for investigating liver physiology and disease modelling, including MASLD. Furthermore, it offers a promising tool for drug discovery, efficacy assessment, and toxicity screening, thereby potentially reducing reliance on preclinical animal models and supporting data for prospective clinical trials.

## Acknowledgement

This work was financially supported by the Indian Council of Medical Research (ICMR) (Grant ID: 2021-12154 and IIRPSG-2025-01-03369), New Delhi, India.

## Materials and Methods

Unless otherwise specified, the chemicals used in the current work were purchased from Thermofisher. All cell culture incubations were maintained in a standard humidified environment at 37°C and 5% CO_2_. The Institutional Ethics Committee [(THS 1.8.1/ (146)] of BRIC-Translational Health Science and Technology Institute, Faridabad, India, approved this study.

### hPSCs culture and maintenance

The hPSCs utilised in this study included the human embryonic stem cell line (hESCs) H1 (WiCell) and human induced pluripotent stem cells (hiPSCs) UCSD002i-16-1 (WiCell) [51]. These cell lines were cultured in either Essential 8 (E8) media or ExCellerate™ iPSC Expansion Media (Bio-Techne), both supplemented with 1% Penicillin-Streptomycin in tissue culture plates (TCP) coated with Matrigel hESC-Qualified Matrix, LDEV-free (Corning)[29]. For passaging, hPSCs were dissociated using 0.5 mM EDTA, plated, and maintained as described above. Cells were passaged every 3 to 4 days (confluency ∼ 70-80%) using 0.5 mM EDTA. Pluripotency was monitored by immunostaining the cells for the pluripotency markers NANAOG and OCT4.

### hPSCs differentiation into definitive endoderm (DE)

For DE differentiation, the initial hPSCs were seeded as an accutase dissociated single cell passaging protocol, as described previously[29]. In the presence of 10μM Y-27632, these dissociated hPSCs were seeded onto Matrigel-coated 6-well tissue culture plates (TCP). After 12-18 h, when the hPSCs reached nearly 40-50% confluency, the cells were treated with differentiation media comprised of RPMI / B-27 Media with insulin, supplemented with 3-4 µM CHIR99021 for 24-27 [28, 29] (day 0). On day 1, cells were subsequently treated with RPMI/B-27 media supplemented with insulin for 24-27h in the absence of CHIR99021. Depending on the type of hPSCs lines, the CHIR99021 concentration and cell number need to be optimised for DE differentiation.

### DE differentiation into hepatoblast (HE)

On day 2, we utilised accutase dissociated single cell passaging with a known cell number of DE in a multiwell cell culture platform, e.g., a 96-well format [22, 26]. The DE cells were dissociated using accutase and seeded at a density of 79,000-80,000 cells/cm² on Matrigel-coated tissue culture plates (TCP) in presence of 10μM Y-27632 for HE specification. The seeding density may require optimisation depending on the specific hPSCs line. The cells were cultured in differentiation media with 1% DMSO, which consisted of Knockout DMEM basal media supplemented with 10% KnockOut Serum Replacement, 1% Glutamine, 1% MEM Non-Essential Amino Acids Solution (NEAA, 100X), and 0.1 mM 2-mercaptoethanol. The media was changed every 24 h, and cells were cultured for an additional 4 days.

### HE differentiation into hepatocyte like cells (HLCs)

On day 7, cells were cultured in differentiation medium with 100 nM dexamethasone to differentiate HE cells into HLCs for three days. After day 10, maintenance medium was used for HLCs maintenance. The maintenance medium is composed of Williams’ E Media basal media supplemented with 10% foetal bovine serum (FBS, BioWest), 10 µM hydrocortisone-21-hemisuccinate (Merck), 1% Insulin-Transferrin-Selenium (100X), 1% GlutaMAX (100X), 100 nM dexamethasone (Merck), and 1% Penicillin-Streptomycin.

### Differentiation of hPSCs to hepatic liver organoids (HLOs)

The hPSCs were differentiated in HLOs, the same 10-day differentiation protocol used for HLCs was applied under three-dimensional culture conditions. Briefly, hPSCs were dissociated into single cells using Accutase and seeded into Corning Elplasia® 96-well Microplate to facilitate uniform spheroid formation. Cells were cultured in the presence of 10μM Y-27632 for the initial 16-18 h to promote cell survival and enable spontaneous aggregation into compact spheroids. Following aggregate formation (day 0), differentiation was initiated with CHIR99021 (3-4 µM) mediated DE induction (day 0–2), identical to the HLCs differentiation. Hepatic specification was then achieved using differentiation medium containing 1% DMSO from day 2 to day 7, followed by differentiation medium containing dexamethasone (100 nM) from day 7 to day 10. Media changes and small molecule treatments were performed as described for hPSC-derived HLCs. By day 10, organoids were harvested for downstream analyses, including morphological assessment, gene expression, immunostaining, and functional assays.

### Quantitative gene expression analysis

The total RNA was isolated from hPSCs (day 0), DE (Day 2), HE (Day 7), and HLCs and HLOs (Day 10) following TRIzol reagent according to the manufacturer’s protocol. The cDNA was synthesised using 500 ng total RNA with iScript™ cDNA Synthesis Kit (Biorad). The gene expression was determined by quantitative RT-PCR on a Biorad real-time PCR system (Biorad) using SYBR green master mix or Taqman FAST Universal PCR Master Mix. Relative gene expression quantitation analysis for the target gene was conducted according to the 2^(ΔΔ^ ^threshold^ ^cycle)^(2^-ΔΔDDCT^) method. All data were normalised to ACTIN (ACTB) or GAPDH. Primer sequences and Taqman probe are listed in Table 2.

**Table 2.**
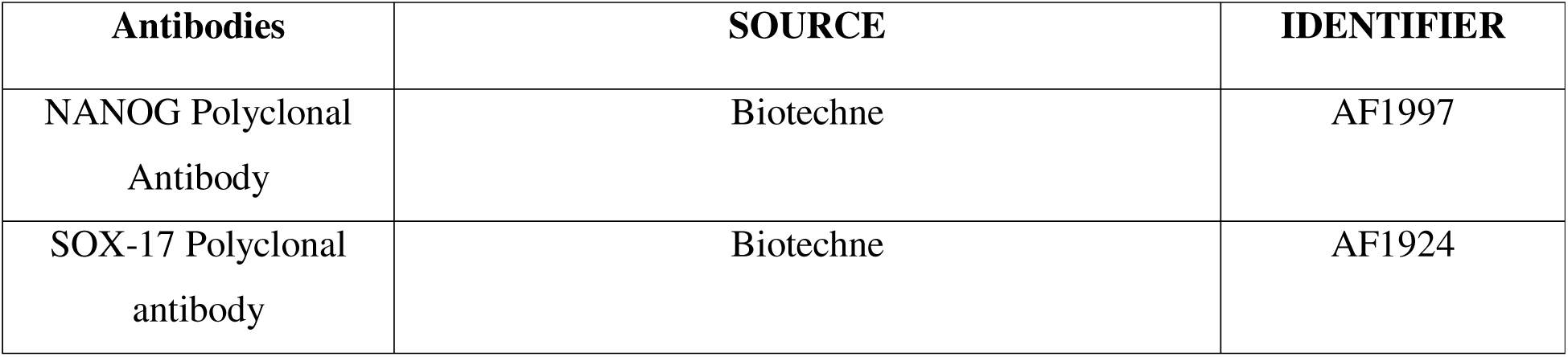
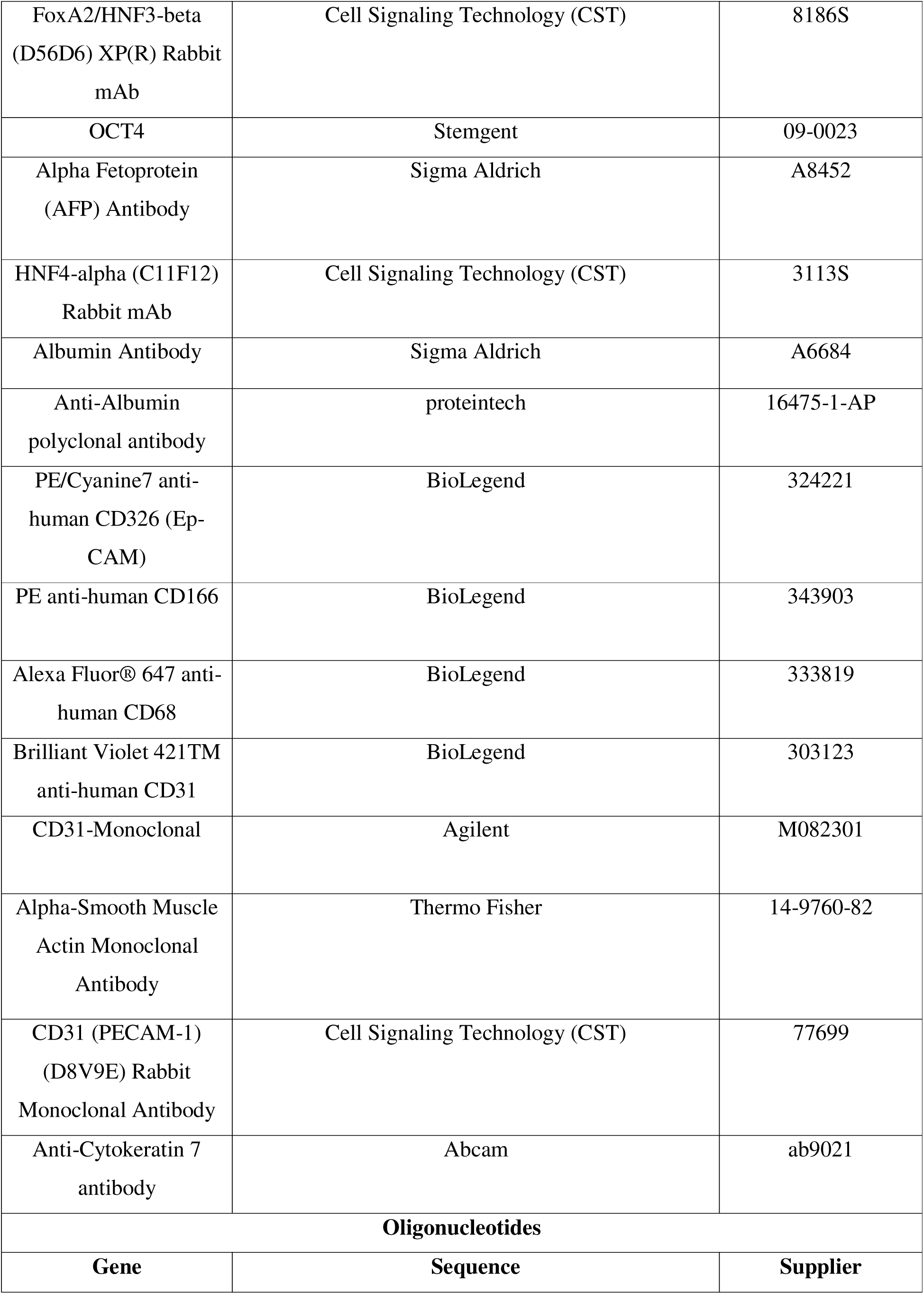
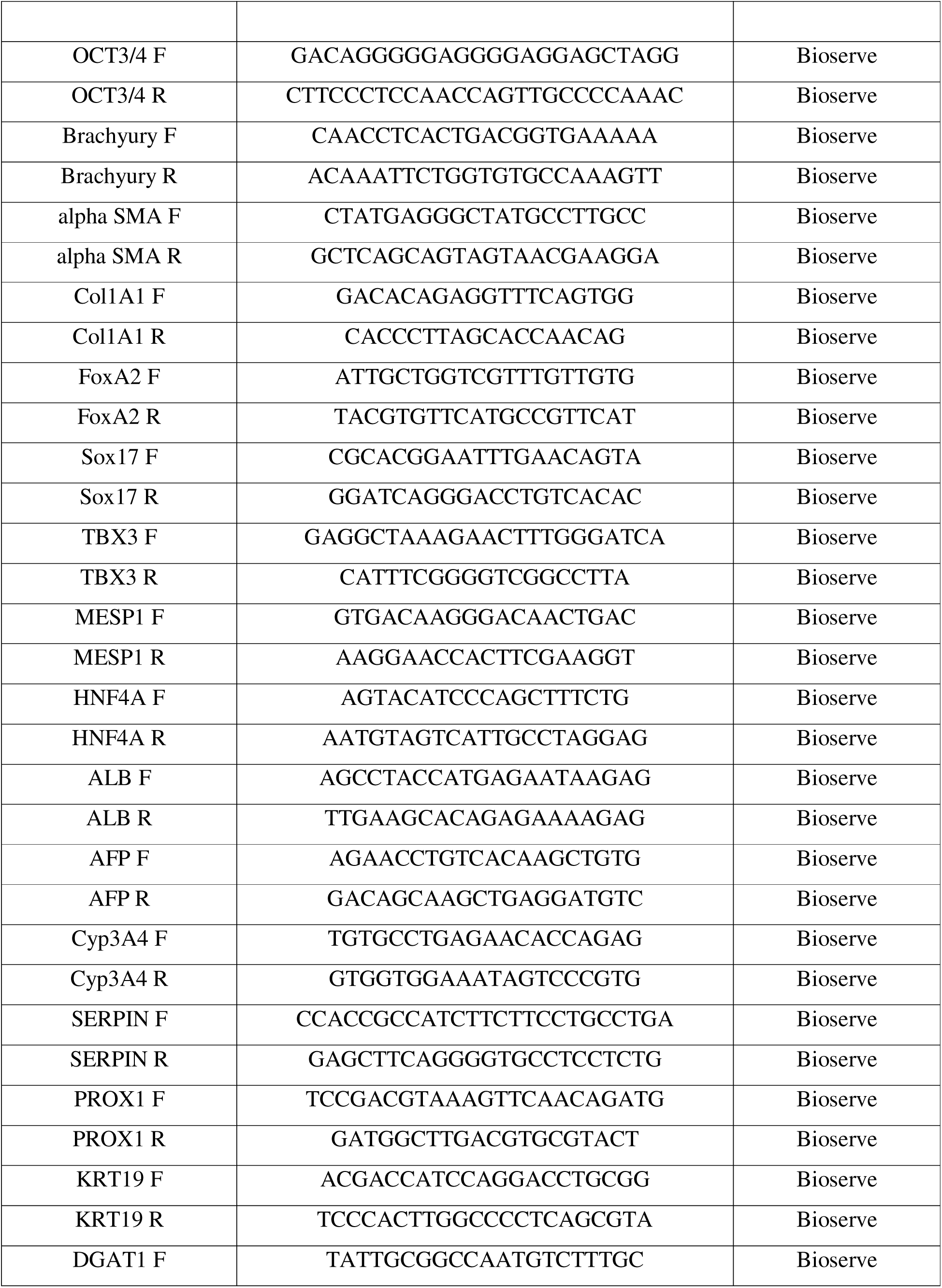
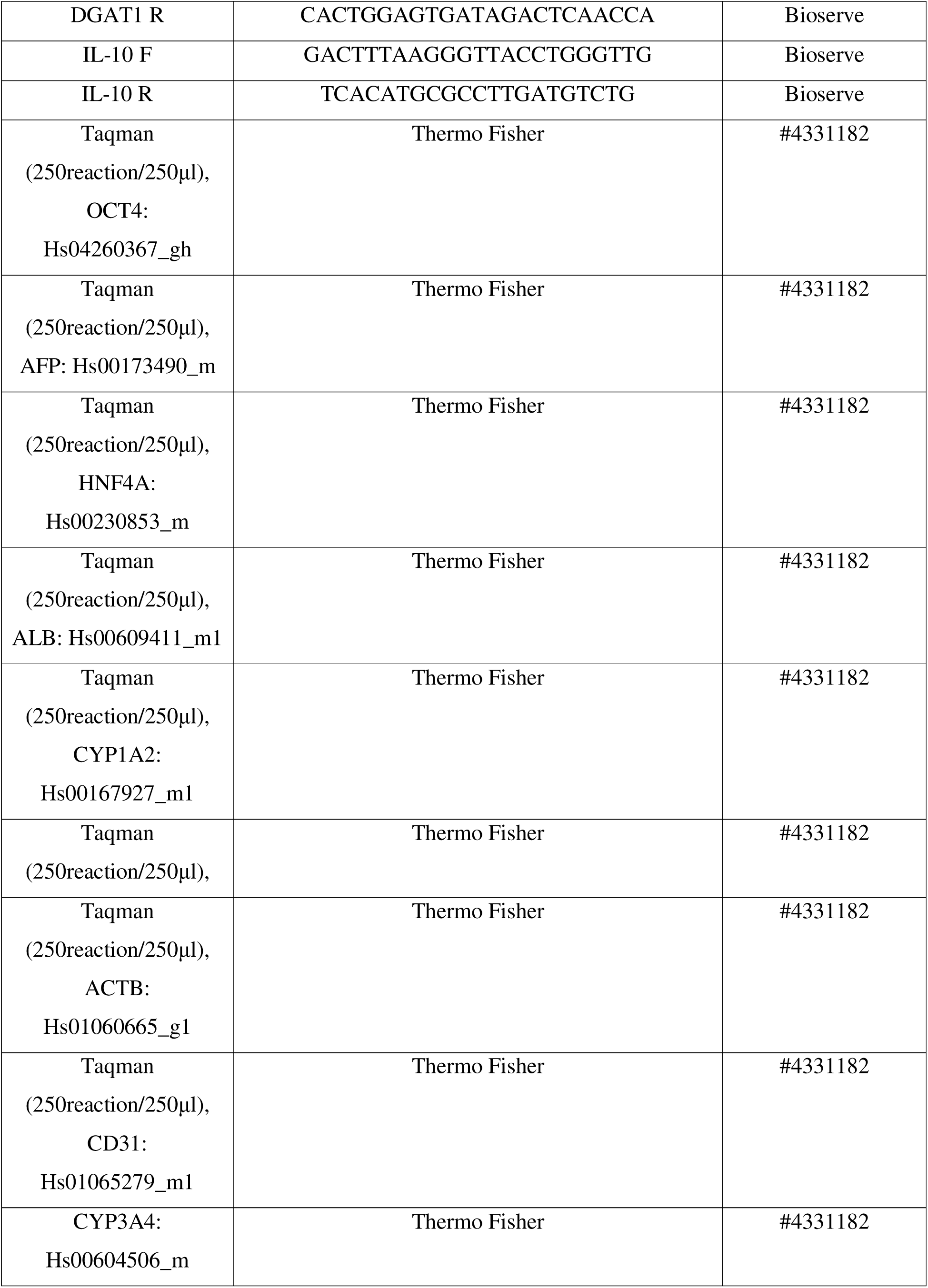
List of antibodies and primer used in the study.

### Protein quantification

The HLCs were lysed with lysis buffer (10 mM Tris, 150 mM NaCl, 0.1% Triton-X100, 1 mM EDTA, 0.1 % SDS, 0.5% sodium deoxycholate with an EDTA-free protease inhibitor cocktail on ice for 10-15 min)[50]. Cell lysates were centrifuged and pelleted down at max speed (12000xg) at 4°C for 15 min to remove cell debris. The supernatant was collected, and protein concentration was measured by Pierce™ BCA Protein Assay Kit according to the manufacturer’s protocol.

### Albumin secretion quantification

Human Serum Albumin DuoSet ELISA (Bio-Techne) was used to measure albumin secretion by HLCs and HLOs in conditioned media on day 10. Day 10 supernatant was collected, and the cells were subjected to protein quantification for normalisation. Briefly, a 96-well plate was coated with capture antibody overnight provided with the kit. The next day, all the regents were prepared according to the manufacturer’s protocol, and the ELISA was performed with samples that were 1:50 diluted. The absorbance was read by spectrophotometer (SpectraMax® i3x Multi-Mode Microplate Reader) at 450 nm.

### Quantification of urea synthesis

On day 10, the HLCs were incubated for 24 h after adding the fresh media. Then, supernatants were collected and analysed to determine the amount of urea production using the QuantiChrom Urea Assay Kit (BioAssay Systems). The assay was performed according to the manufacturer’s instructions. The amount of urea secretion was calculated according to each standard, followed by normalisation of the total protein content.

### Flow cytometry

Expression of DE markers were assessed by flow cytometry on day 2. The DE was dissociated with Accutase. For day 10 cells of HLOs and HLCs, a sequential treatment of collagenase type IV (∼30 min) followed by Accutase (∼30 min) was used. Cells were fixed with 1% Paraformaldehyde (PFA) for 10 min at room temperature (RT), then permeabilised with staining buffer (2% FBS +0.1% triton x100 in PBS), cells were incubated with primary antibody (diluted in staining buffer) for 1 h at 4°C, followed by three washes. Secondary antibody incubation was performed for 30 min at 4°C, with three subsequent washes. Cells were suspended in PBS for flow cytometry analysis (FACS CANTO II analyser, BD Biosciences) and analysed with FlowJo software .

For neutral lipid quantification, sHLCs were dissociated with collagenase and accutase as mentioned above and fixed with 1% PFA for 10 min. After washing with PBS, the cells were stained with Nile Red for 10 min at RT. After incubation, the cells were washed 3-4 times with PBS and immediately used for flow cytometry analysis.

### CYP3A4 assay

To measure CYP3A4 induction in HLCs, the gene expression levels of CYP3A4 were measured by quantitative gene expression analyses performed with TaqMan Gene Expression Assays ( Primers listed in Table 2). Briefly, the HLCs were treated with 50 µM dexamethasone for 48h, which is known to induce CYP3A4. The relative quantification of target gene expression levels was performed using the 2^-ΔΔCT^ method. Values were normalised to those of the housekeeping gene ACTIN (ACTB).

### CYP3A4 activity assay

For HLOs to be tested for CYP3A4 activity, 50 µM dexamethasone was supplemented to the differentiation media and administered to the cells twice for a duration of 48 h via daily media change. CYP3A4 activity was determined with the P450-Glo™ CYP3A4 Assay (Promega, WI, USA) and normalized by cell viability, which was determined with the CellTiter-Glo® Luminescent Cell Viability Assay (Promega). Both experiments were carried out according to the manufacturer’s protocol.

### Preparation of free fatty acids (FFA) stock and treatment

The fatty acids oleic acid (OA) and palmitic acid (PA) were conjugated with fatty-acid-free bovine serum albumin (Capricon, BSA) to prepare free fatty acids (FFA) stocks, as previously described [52]. Briefly, the OA and PA were dissolved in absolute ethanol and chloroform, respectively, to a final concentration of 1 M. The liquid form of fatty acids was dissolved in 0.01 N NaOH in a water bath individually at 70°C for 90 min to yield a final concentration of 20 mM. The 20% BSA was solubilised in PBS and mixed with the liquid form of fatty acids solution in a 1:1 ratio to get an individual 10 mM concentration of each fatty acid. The prepared mixture was vortexed and then incubated for 1 h at 42°C for conjugation. After cooling to RT, the mixture was sterile filtered using a 0.2 μm filter. These conjugated fatty acids were stored at −20°C. For FFA treatment to generate sHLCs and sHLOs, HLCs or HLOs were cultured in maintenance media and a combination of FFA at a ratio of 2:1 (OA: PA) and final concentrations of 0.2, 0.4, 0.8, 1.6, 3.2, and 6.4 mM for 48 h and 120 h. The BSA alone served as a vehicle.

To establish a curative model of MASLD, HLCs and HLOs were first induced into a steatotic state by treatment with FFA for 120 h. After steatosis confirmation, sHLCs and sHLOs were treated with Resmetirom (Medchem) at a concentration of 100 µM in the continued presence of FFA for an additional 120 h. Vehicle-treated steatotic cells served as controls. Post-treatment, cells were harvested for assessment of intracellular TGs levels, lipid accumulation (BODIPY staining), and gene expression analysis of lipogenic (DGAT1, DGAT2), inflammatory (IL-10), and fibrotic (COL1A1, alpha SMA) markers for steatosis or steatohepatitis.

### Oil Red O (ORO) staining and quantification

The sHLCs were washed three times with PBS and fixed with 4% PFA. Following fixation, cells were washed twice with distilled water and treated with 60% isopropanol for 5 minutes, then allowed to air-dry completely at room temperature. A fresh ORO working solution was prepared by mixing 6 ml of ORO stock solution (0.35 g Oil Red O dissolved in 100 ml isopropanol, filtered through 0.2 µm) with 4 ml of distilled water, allowed to equilibrate for 20 minutes at room temperature, and filtered through a 0.2 µm filter prior to use. The working solution was applied to cells for 10 minutes at room temperature, followed by four washes with distilled water. For semi-quantitative analysis, ORO dye was eluted by adding 100% isopropanol with gentle shaking for 10 minutes. The absorbance was read by a spectrophotometer at 500 nm [53].

### Periodic Acid-Schiff (PAS) staining for glycogen

For Periodic Acid-Schiff (PAS) staining, HLCs were fixed with 4% PFA at RT for 20 min and then washed with distilled Milli-Q water (DI). The 0.5% periodic acid solution (Merck) was added to HLCs (50ul in a 96-well plate) for 5 min at RT, followed by washing with DI three times. After washing, Schiff’s solution was added for 15 min at RT and washed with DI three times. The inverted microscope (Nikon, Japan) was used to capture bright-field micrographs.

### Immunostaining

To perform immunostaining on cells representing various differentiation stages, the cells were initially fixed with chilled 100% methanol (−20°C) for 15 min at -20°C. The methanol was removed and cells were washed with PBS 3 times. For blocking, cells were incubated in a blocking buffer solution (1% bovine serum albumin in PBS) for 15 min. Primary antibody incubation (in blocking buffer solution) was followed for 1 h. Cells were then washed with blocking buffer solution and incubated with secondary antibodies (in blocking buffer) for 1 h at RT. Nuclear staining was performed with Hoechst 33342, and images were acquired using the Olympus IX83 inverted microscope (Olympus, Japan).

### H2DCFDA assay

Intracellular ROS formation was measured using the fluorogenic substance dichlorodihydrofluorescein diacetate (H2DCFDA) according to the manufacturer’s protocol. For ROS measurement, the culture media of sHLCs was replaced with advanced serum free DMEM containing 10 µM H2DCFDA, followed by incubation for 30 min at 37 °C, 5% CO2 in a humidified incubator. Subsequently, the supernatants were aspirated, and the cells were dissociated with collagenase IV for 30 min, followed by accutase for 30 min. After dissociation, cells were washed with PBS and analysed immediately using flow cytometry.

### BODIPY staining

The sHLCs or sHLOs were washed twice with PBS and fixed in 4% PFA for 10 min. Following fixation, the cells were washed twice with PBS and stained with 5 μg/ml BODIPY for lipid detection and Hoechst 33342 for nuclear staining. After a 30 min incubation at RT, the cells were washed twice with PBS. Confocal images were captured using a confocal laser scanning microscope (Olympus FV3000, Japan).

### TGs quantification

The sHLCs or sHLOs were washed with PBS, resuspended in a 5% NP-40/ddH2O solution, and collected in a microcentrifuge tube. The samples were then slowly heated to 85°C in a water bath for 2–5 min, or until the solution became cloudy, and then allowed to cool on ice. This heating and cooling process was repeated for 3-4 times. Then the samples were centrifuged for 10 min at 12000xg in a microcentrifuge to remove any insoluble material. The levels of TG were measured in the supernatants using the Randox TRIGS kits according to the manufacturer’s protocol [54].

### Metabolomics

On day 10, HLCs were treated with FFA (800µM) for 48 h with every 24 h media renewal. For control, cells were treated with BSA alone. After 48 h, cells were dissociated with collagenase type IV (30 min) and then accutase (30 min). The dissociated cells were subjected to metabolomics analysis [55]. Briefly, metabolite analysis was performed using an Orbitrap Fusion mass spectrometer equipped with a heated electrospray ionization (HESI) source. Data were acquired in both positive and negative ionization modes across a mass range of 60-900 Da. MS1 data were collected at a resolution of 120,000, while MS2 data were acquired at a resolution of 30,000. Chromatographic separation was achieved using an Ultimate 3000 UPLC system with both hydrophilic interaction liquid chromatography (HILIC) and reverse phase (RP) columns. HILIC separations were performed on an XBridge BEH Amide column (Waters Corporation) with a mobile phase consisting of (A) 20 mM ammonium acetate in water (pH 9.0) and (B) 100% acetonitrile. A gradient elution from 85% B to 10% B over 14 minutes at a flow rate of 0.35 ml/min was employed. RP separations utilised an HSS T3 column (Waters Corporation) with (A) water and (B) methanol, both containing 0.1% formic acid. A gradient elution from 1% B to 95% B over 10 min at a flow rate of 0.3 ml/min was used. For both HILIC and RP separations, the injection volume was 5 μl.

### Preclinical study

Animal experiments were conducted in accordance with protocols approved by the Institutional Animal Ethics Committee (IAEC/THSTI/326) and the guidelines of the Committee for Control and Supervision of Experiments on Animals (CCSEA), Government of India. Male and female NOD SCID mice (6-7 week, n=6) (The Jackson Laboratory) were housed with a 12-h light–dark cycle and ad libitum access to water and standard rodent diet. The transplantation of the organoids under the kidney capsule or the sham laparotomy was performed as described in [56] with some modifications. In brief, anesthesia was induced intraperitoneal administration of ketamine (80 mg/kg) combined with xylazine (10 mg/kg). Preoperative analgesia was provided by subcutaneous injection of buprenorphine (0.05 mg/kg). Following a sterile preparation of the left flank, a 1.5□cm incision was made midway between the last rib and the iliac crest and approximately 0.5□cm parallel and ventral to the spine. The left kidney was slowly externalized through the abdominal incision using sterile cotton swabs, immobilized using nontraumatic forceps, and moisturized with warm sterile saline. The injection site was located at the upper lateral side of the kidney, and a 1□ml syringe with a Venflon IV Cannula 22G containing either the organoid suspension or pure Matrigel (Corning) (sham surgery) was gently pushed under the capsule toward the inferior pole of the kidney to avoid perforation and damage to the blood vessels. The 15-20 µl of organoid suspension or Matrigel matrix was very slowly injected beneath the kidney capsule, and the needle was simultaneously slowly withdrawn from the capsule to avoid backflow. Next, the kidney was placed back in the body cavity, and the abdominal wall was closed with sutures (6-0 Prolene). After surgery, the mice were examined daily during the first week for normal wound healing, with weight measurements; thereafter, once a week. Blood was collected once a week.

Animals were euthanized using CO_2_ asphyxiation at week 4, and left kidney tissue was immediately harvested and rapidly frozen in liquid nitrogen, then stored at −80°C until processing. For immunostaining, frozen tissue samples were transferred from −80°C storage and embedded in Optimal Cutting Temperature (OCT) medium for cryosectioning. Serial cryosections (10 μm thickness) were cut using a cryostat (Leica) and mounted onto poly-L-lysine-coated glass slides. Immunostaining was performed using a standardized protocol-slides were equilibrated at room temperature for 1–2 minutes, washed three times with 1X PBS for 5 minutes each, fixed with 4% paraformaldehyde for 15 minutes, and washed three times with 1X PBS with Triton X-100 (PBST; 0.1% Triton X). Antigen retrieval was performed by dipping slides into preheated sodium citrate buffer (880 mg in 300 mL milliQ water, pH 6.0) for 10 minutes in a microwave with intermittent boiling, followed by cooling to 4°C for 30–45 minutes and three additional washes with 1X PBST. Non-specific binding was blocked using 3% bovine serum albumin (BSA) in 1X PBST in a humidified chamber for 2-3 h, followed by incubation with primary antibodies (diluted in 1X PBST) in a humidified chamber at 4°C overnight. On next day, slides were washed three times with 1X PBST for 5 minutes each, incubated with species-appropriate fluorescent-conjugated secondary antibodies (diluted in 1X PBST) in a humidified chamber at 4°C for 2 h, washed three additional times with 1X PBST, and mounted using mounting medium. Confocal images were captured using a confocal laser scanning microscope (Olympus FV3000, Japan).

### Statistical analysis

All data analyses and statistical tests were performed using GraphPad Prism software (version 9.0). Results were expressed as mean ± SD. Statistical significance were evaluated using one-way or two-way ANOVA, followed by Tukey’s post hoc test. Where n.s., not significant, *p < 0.05, **p > 0.01,***p > 0.001, ****p > 0.0001.

